# TROP2 and immune gene expression by African ancestry in triple-negative breast cancer

**DOI:** 10.64898/2026.10.29.762976

**Authors:** Alvan Okechukwu, Emmanuel O. Akala

## Abstract

**Purpose:** TROP2 is a treatment target in triple-negative breast cancer (TNBC). *TACSTD2* encodes TROP2, while *PDCD1* encodes the immune protein PD-1. We investigated whether these genes differ by race or African genetic ancestry and whether the reported link between higher *TACSTD2* and lower immune-gene expression partly reflects the mixture of cells in tumor samples.

**Methods:** We analyzed TNBC tumors from TCGA (n = 107) and GSE268851 (n = 253; ancestry data for 229), together with spatial, normal-breast, and protein datasets. Regression models adjusted for age, tumor stage, and a score based on epithelial gene expression. We compared *TACSTD2* with other epithelial markers using paired bootstrap analysis.

**Results:** *TACSTD2* was detected in nearly all tumors, with no significant differences by race or ancestry. Higher African ancestry was linked to higher *PDCD1* expression in TCGA (P = 3.9 × 10^−4^), but not in GSE268851. In GSE268851, higher *TACSTD2* remained associated with lower *CD8A, GZMB*, and cytotoxic score after epithelial adjustment (all q < 0.003). These associations were stronger than those for *EPCAM*; the *CD8A* association was also stronger than that for *KRT8. TACSTD2* was 1.55-fold higher in TNBC than in normal breast and correlated with TROP2 protein (ρ = 0.74).

**Conclusion:** *TACSTD2* was widely expressed in the TNBC groups studied, while the ancestry–*PDCD1* association differed between cohorts. Epithelial gene expression did not fully explain some negative *TACSTD2*–immune associations, and some were stronger than those for other epithelial markers such as *EPCAM*.

## 1. Introduction

Triple-negative breast cancer (TNBC) lacks estrogen receptor, progesterone receptor, and HER2 expression. Black women experience a greater burden of TNBC than White women [1–3]. Race and genetic ancestry are not the same. Race reflects how people are classified in society and the experiences linked to those classifications. Genetic ancestry uses DNA to estimate inherited population background [4]. Examining both may help us understand how tumor biology and treatment targets differ between populations.

Two important treatment targets in TNBC are TROP2 and PD-1. TROP2 is a protein found on many breast cancer cells and is encoded by *TACSTD2*. Sacituzumab govitecan binds TROP2 and delivers the cancer-killing drug SN-38 [5, 6]. PD-1, encoded by *PDCD1*, is found mainly on immune cells. Pembrolizumab blocks PD-1 to help immune cells attack cancer [7, 8]. In June 2026, the US Food and Drug Administration approved sacituzumab govitecan alone for certain patients with advanced TNBC who cannot receive PD-1/PD-L1 inhibitors, and with pembrolizumab for previously untreated PD-L1-positive advanced TNBC (combined positive score [CPS] ≥10) [9]. In ASCENT-04/KEYNOTE-D19, the combination prolonged median progression-free survival compared with chemotherapy plus pembrolizumab (11.2 vs 7.8 months; hazard ratio, 0.65) [10]. Other TROP2-directed treatments are also being studied [11–15].

Studies have reported differences in immune-related gene expression among TNBC populations, but the findings have not always agreed [16–18]. Martini et al. found that African ancestry was associated with *PDCD1* and other immune genes [3]. It remains unclear whether this result can be reproduced in other patient groups or whether *TACSTD2* expression differs by race or genetic ancestry. Previous studies found that breast tumors with higher TROP2 expression often have lower expression of immune-related genes [19–21]. Wu et al. also showed experimentally that TROP2 can help prevent immune cells from entering breast tumors, through a process involving claudin proteins [22]. This relationship is not easy to interpret from ordinary RNA sequencing. A tumor sample contains a mixture of cancer cells, immune cells, and other cells. If one tumor contains more cancer cells and fewer immune cells, it may show higher *TACSTD2* and lower immune-gene expression even without a direct effect of TROP2 on immune cells. We therefore adjusted for a score based on epithelial genes and compared *TACSTD2* with other epithelial markers. This score helps account for epithelial-related differences between samples, but it does not directly measure the number of cancer cells.

We first tested whether *TACSTD2* and *PDCD1* expression differed by race or African ancestry in independent TNBC cohorts. We then examined whether negative *TACSTD2*–immune associations remained after adjusting for epithelial gene expression and whether they were stronger than associations involving *EPCAM, KRT8*, or *KRT18*. Spatial, normal-breast, and protein data helped us interpret the results.

## 2. Materials and Methods

### 2.1 Study design and cohorts

We analyzed two public TNBC datasets (Table 1). TCGA-BRCA included 107 ER-, PR-, and HER2-negative primary tumors with ancestry estimates from Carrot-Zhang et al. [4]. GSE268851 included 253 Louisiana TNBC tumors (128 Black and 125 White women); 229 had ancestry data [23]. GSE268851 used capture-based RNA sequencing of formalin-fixed surgical tissue containing at least 50% tumor, whereas TCGA used fresh-frozen tissue and poly(A)-selected sequencing.

**Table 1.** Datasets and their roles in the analysis.

| **Dataset** | **Sample size** | **Material** | **Main analysis** |
|---|---|---|---|
| GSE268851 | 253 TNBC; 229 ancestry-typed | FFPE bulk RNA-seq | Race, ancestry, epithelial-immune associations |
| TCGA-BRCA | 107 TNBC | Bulk RNA-seq | Ancestry and immune expression |
| GeoMx (Zhu et al.) | 57 patients | Spatial RNA | Epithelial versus immune expression |
| TCGA-GTEx Toil | 107 TNBC; 80 normal | Bulk RNA-seq | Tumor versus normal breast |
| GSE205725 | 190 donors; 170 compared | Healthy breast RNA | Expression by race |
| CPTAC breast | 122 tumors; 28 TNBC | RNA and proteomics | RNA-protein correlation |

FFPE, formalin-fixed paraffin-embedded; TNBC, triple-negative breast cancer. The TCGA tumors in the Toil analysis overlap with those in the primary TCGA cohort and do not constitute an independent validation dataset. Genetic ancestry was available for 229 GSE268851 tumors; race was analyzed separately.

To find where *TACSTD2* was expressed within tumors, we used spatial RNA data from Zhu et al. [24] (Zenodo 12752405): 57 patients and 294 sampled regions. We compared each patient’s median log2-transformed, Q3-normalized expression in PanCK-positive epithelial and CD45-positive immune regions using paired Wilcoxon tests. Racial comparisons used Welch tests.

To compare tumors with normal breast, we used uniformly processed TCGA-TARGET-GTEx data from UCSC Xena [25]: 107 TNBC tumors and 80 normal female breast samples. We converted log2(TPM + 0.001) values to TPM and compared ratios of median expression, using 10,000 bootstrap resamples for 95% confidence intervals and Mann–Whitney tests for the distributions. The unpaired samples differ in cell composition, so this does not directly measure increased expression within cancer cells [26].

Healthy-breast GSE205725 data included 170 donors (62 African American, 108 White) from 190 available [27]. Mann– Whitney tests compared CPM values; *PDCD1* detection was evaluated at >0 and >1 CPM.

CPTAC data [28] included 122 breast tumors (28 TNBC). We assessed *TACSTD2* mRNA (log2 FPKM) versus total TROP2 protein (log2 abundance ratio) using Spearman correlation and approximate 95% confidence intervals from Fisher’s z-transform.

### 2.2. Expression and ancestry definitions

We used edgeR [29] to convert the full RNA-sequencing count matrices to log2 counts per million (log2CPM; prior count 0.5). The immune panel (Fig. 2c) contains 19 measures: four T-cell genes (*CD3D, CD3E, CD2, TRAC*); six checkpoint or exhaustion-related genes (*PDCD1, LAG3, HAVCR2, TIGIT, CTLA4, TOX*); three cytotoxic genes (*CD8A, GZMB, PRF1*); two regulatory T-cell genes (*FOXP3, IL2RA*); three combined gene-expression scores; and a separate *PDCD1* model adjusted for *CD3E*. Expression of *PDCD1* alone was not taken as proof of T-cell exhaustion.

We calculated gene-expression scores by standardizing each gene as a z-score and averaging the values. We used all 253 GSE268851 or 107 TCGA tumors for standardization, including in subgroup analyses. The epithelial score included *EPCAM, KRT8, KRT18*, and *CDH1* but excluded *TACSTD2*. The cytotoxic score contained *CD8A, GZMB*, and *PRF1*; the exhaustion score contained *LAG3, HAVCR2, TIGIT*, and *TOX*; and the regulatory T-cell score contained *FOXP3, IL2RA*, and *IKZF2*. These scores reflect expression, not immune-cell numbers or activity [30].

Race was analyzed separately from African genetic ancestry. TCGA used continental ancestry proportions from Carrot-Zhang et al. [4]; GSE268851 used the C_W_African component estimated with supervised STRUCTURE from 161 ancestry-informative markers [23]. Values ranged from 0 to 1 but were derived differently. GSE268851 ancestry models included 229 patients, with a Black-only analysis of 118.

In TCGA, low ancestry was defined as <0.05 (n = 73) and high ancestry as >0.595 (n = 27); seven intermediate patients were excluded only from this grouped comparison. Because ancestry estimates clustered near the extremes, continuous-model slopes should not be assumed constant across the entire ancestry range.

### 2.3 Statistical analysis

We used ordinary least-squares regression to test gene-expression associations with race, ancestry, or epithelial markers. HC3 robust standard errors and t-based 95% confidence intervals accounted for unequal variability across samples. Main models adjusted for age, tumor stage, and epithelial score. Direct epithelial-marker comparisons adjusted for age and stage without the epithelial score, because that score includes the comparison genes.

Benjamini–Hochberg correction was applied separately within prespecified families: five SN-38 genes per model; four epithelial markers per immune outcome; 19 TCGA immune measures; five epithelial-adjusted immune outcomes; and 15 paired marker contrasts. Adjusted P values are denoted q.

In GSE268851, we used 10,000 paired patient-level bootstrap resamples (seed 20261021) to compare standardized *TACSTD2* coefficients with those for *EPCAM, KRT8*, and *KRT18*. Both coefficients in a comparison used the same resampled patients. We subtracted the comparator coefficient from the *TACSTD2* coefficient; a negative difference indicated a stronger negative *TACSTD2* association. We calculated percentile 95% confidence intervals and approximate two-sided P values from centered bootstrap distributions, then corrected the 15 comparisons for multiple testing.

For the TCGA ancestry–*PDCD1* association, sensitivity analyses replaced the epithelial score with tumor purity, removed influential observations or individual patients, compared ancestry groups, adjusted for *CD3E*, and examined BMI and recorded chemotherapy where available. Treatment timing was unknown, so recorded chemotherapy was not classified as neoadjuvant.

Analyses were exploratory, without prospective power or equivalence testing. Applicable REMARK reporting elements [31], inclusion lists, software versions, checksums, and detailed outputs are documented in the Supplementary Materials.

## 3. Results

### 3.1 TACSTD2 is widely expressed in TNBC with no significant differences by race or African ancestry

*TACSTD2* was widely expressed in TNBC tumors in both datasets. Expression exceeded 1 log2CPM in 97.6% of GSE268851 tumors and 99.1% of TCGA tumors, with median expression levels of 5.56 and 8.54 log2CPM, respectively.

In GSE268851, *TACSTD2* expression did not differ significantly between Black and White women after adjusting for age, tumor stage, and epithelial gene expression (β = 0.0065, 95% CI: −0.311 to 0.324; P = 0.97; n = 253; **Fig. 1a**). Similarly, African genetic ancestry was not significantly associated with *TACSTD2* expression in the full ancestry-typed cohort (β = −0.153, P = 0.50; n = 229) or among Black women alone (β = −0.644, P = 0.17; n = 118). TCGA showed similar findings, with no significant association between *TACSTD2* expression and continuous African ancestry (β = −0.099, P = 0.80; n = 107) or between the high- and low-ancestry groups (β = −0.079, P = 0.81; n = 100; **Fig. 1b**). Additional analyses adjusting for BMI and examining patient subgroups in GSE268851 produced similar results (Supplementary Table S5).

**Fig. 1.**
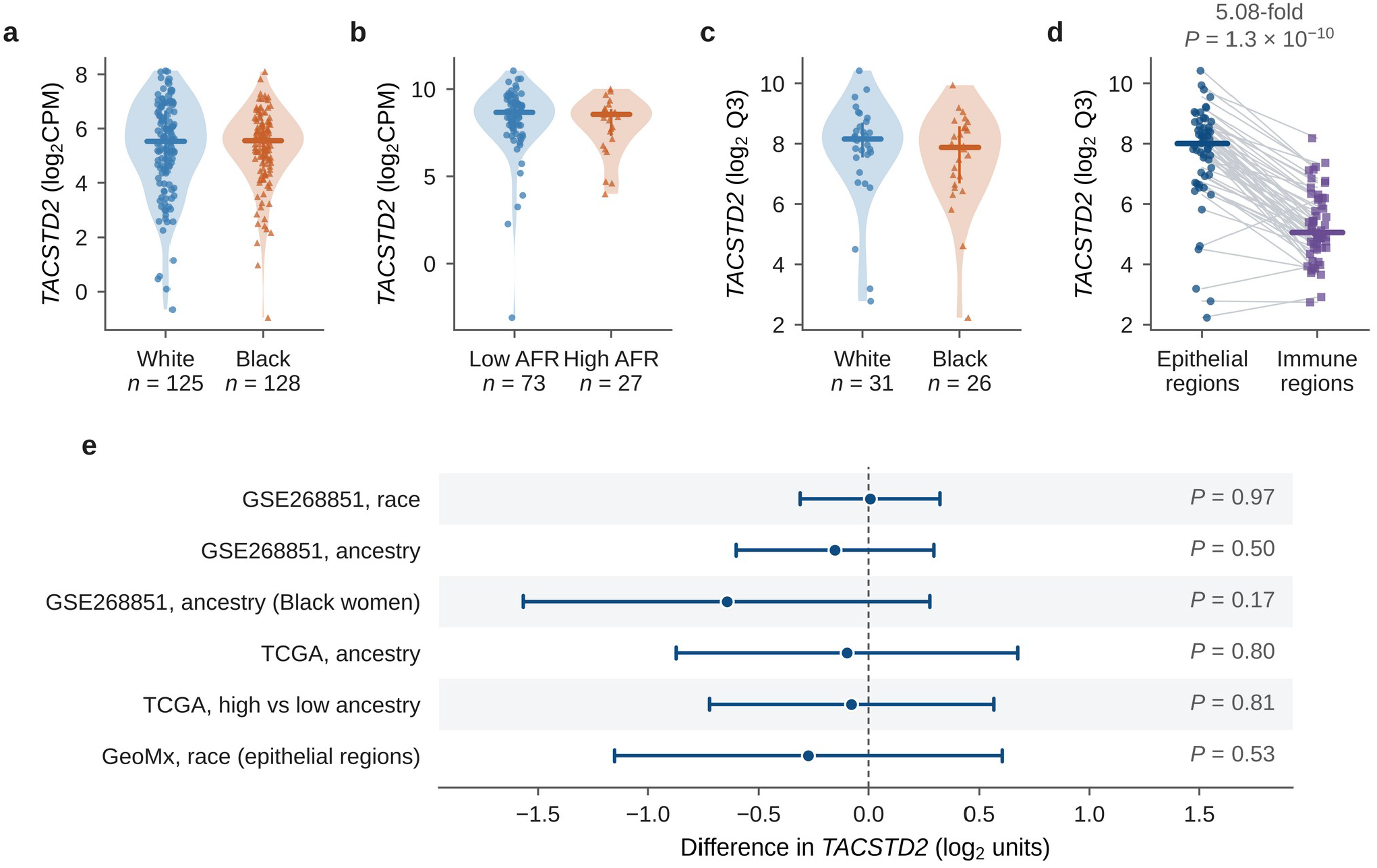
*TACSTD2* expression by race, African ancestry, and tumor region in TNBC. (a) *TACSTD2* expression in tumors from White (n = 125) and Black (n = 128) women in GSE268851. (b) *TACSTD2* expression in TCGA tumors with low (n = 73) and high (n = 27) African ancestry. (c) *TACSTD2* expression in epithelial regions of tumors from White (n = 31) and Black (n = 26) women in the GeoMx spatial dataset. (d) Paired comparison of *TACSTD2* expression in epithelial and immune regions from 57 patients; lines connect regions from the same patient. (e) Race and ancestry associations with *TACSTD2* expression across GSE268851, TCGA, and GeoMx. In panels a–c, circles show White women or low-ancestry tumors and triangles show Black women or high-ancestry tumors; in panel d, circles show epithelial regions and squares show immune regions. Violin plots show expression distributions, points show individual patients, horizontal bars show medians, and vertical lines show interquartile ranges. In panel e, points show regression estimates for the bulk cohorts and the unadjusted mean difference for GeoMx; horizontal lines show 95% confidence intervals, and the dashed line marks no difference. Bulk models were adjusted for age, tumor stage, and epithelial score; the GeoMx comparison used an unadjusted Welch test. Each coefficient represents the labeled race contrast or the change across the 0-to-1 ancestry scale, so values are not directly comparable between rows

**Fig. 2.**
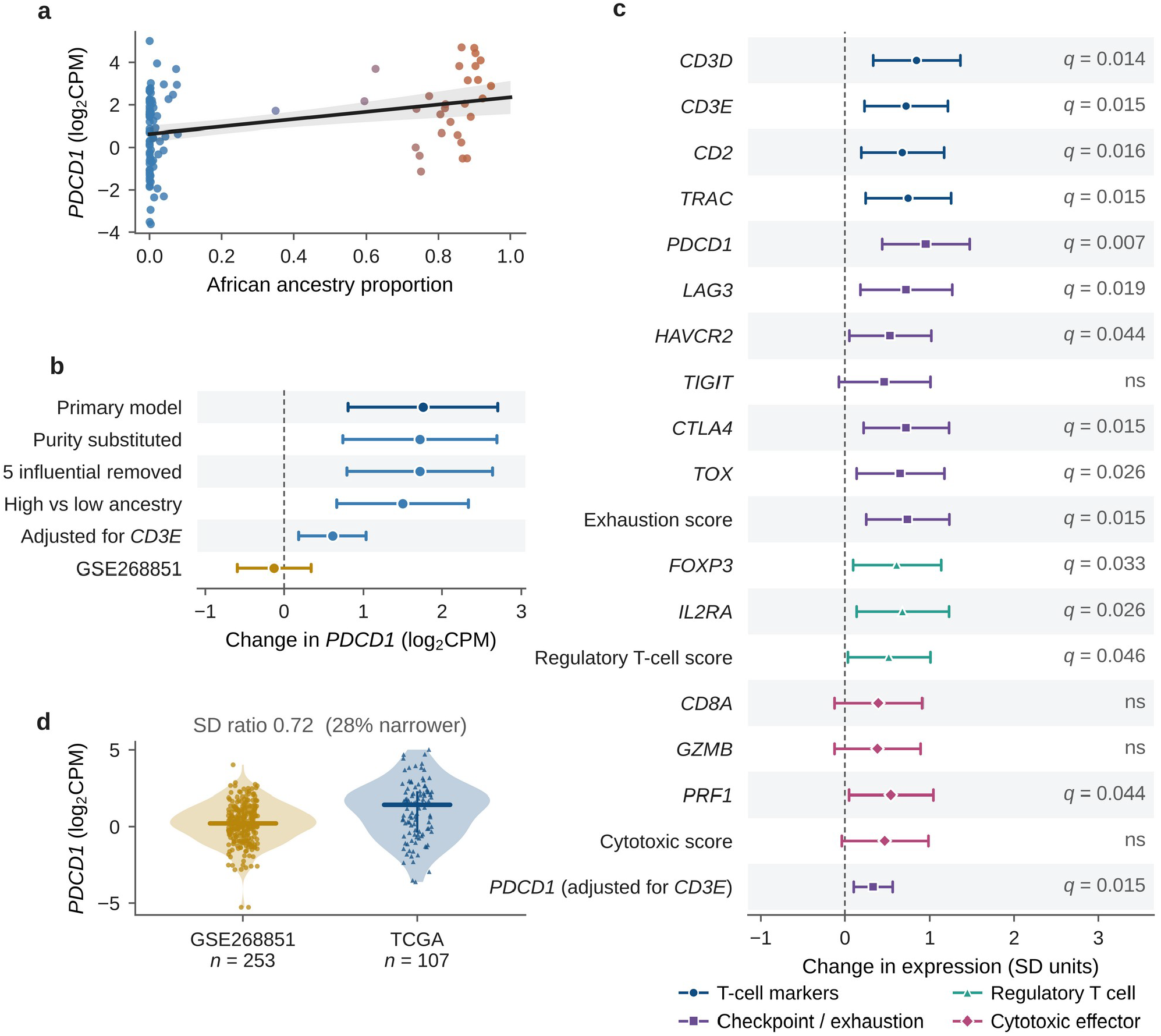
Association of African ancestry with *PDCD1* and immune-related gene expression in TNBC. (a) *PDCD1* expression versus African ancestry proportion in TCGA (n = 107). Points show individual patients, and the line shows an unadjusted linear fit with its 95% confidence band. (b) Adjusted association between African ancestry and *PDCD1* expression in TCGA in the primary model and sensitivity analyses, with the corresponding GSE268851 estimate for comparison. (c) Associations between African ancestry and 19 immune-related measures in TCGA. Marker shapes indicate gene groups: circles, T-cell markers; squares, checkpoint or exhaustion; triangles, regulatory T cell; diamonds, cytotoxic effector. q values are Benjamini–Hochberg-adjusted across the 19 tests; ns, not significant after correction. (d) Distribution of *PDCD1* expression in GSE268851 (circles; n = 253) and TCGA (triangles; n = 107); horizontal bars show medians. The standard deviation of *PDCD1* expression was 28% lower in GSE268851 than in TCGA (SD ratio = 0.72). In panels b and c, points show regression coefficients, horizontal lines show 95% confidence intervals, and dashed lines mark no association. Models were adjusted for age, tumor stage, and epithelial score unless otherwise indicated

We also examined where *TACSTD2* was expressed within TNBC tumors using spatial transcriptomic data. Expression in epithelial regions did not differ significantly between tumors from Black and White women (P = 0.53; **Fig. 1c**).

However, *TACSTD2* expression was approximately 5.08 times higher in epithelial regions than in immune regions (P = 1.3 × 10^−10^; **Fig. 1d**). Higher expression in epithelial regions was observed in 54 of 57 patients.

Overall, *TACSTD2* was widely expressed in TNBC and showed substantially higher expression in epithelial than immune regions. We found no statistically significant differences in its expression by race or African genetic ancestry across the studied cohorts (Fig. 1e).

### 3.2 Higher African ancestry is associated with increased PDCD1 expression in TCGA but not GSE268851

In TCGA, higher African genetic ancestry was significantly associated with higher *PDCD1* expression after adjusting for age, tumor stage, and epithelial gene expression (β = 1.757, 95% CI: 0.807 to 2.707; P = 3.9 × 10^−4^; n = 107; **Fig. 2a**). This relationship remained significant in several additional analyses. Similar results were obtained when tumor purity replaced the epithelial score (β = 1.718, P = 7.2 × 10^−4^), after excluding five influential patients (β = 1.717, P = 3.8 × 10^−4^), and when comparing patients with high versus low African ancestry (β = 1.498, P = 5.8 × 10^−4^; n = 100). The association also remained significant when patients were removed one at a time. After adjusting for *CD3E*, a gene associated with T cells, the ancestry coefficient decreased by approximately 65% but remained significant (β = 0.610, P = 0.0054; **Fig. 2b**). This suggests that differences in T-cell-related gene expression partly account for the observed relationship between African ancestry and *PDCD1* expression.

We also examined whether African ancestry was associated with other immune-related genes in TCGA. Of the 19 immune measures analyzed, 15 remained significant after correction for multiple testing (q < 0.05; **Fig. 2c**). These included all four T-cell markers, seven of eight checkpoint and exhaustion-related measures, all three regulatory T-cell measures, and *PRF1*. However, *CD8A, GZMB*, and the cytotoxic score were not significant after correction. These findings show that higher African ancestry was associated with broader changes in T-cell-related and immune-regulatory gene expression, but not with a significant increase in the overall cytotoxic score.

In contrast, we did not observe a significant association between African ancestry and *PDCD1* expression in GSE268851 (β = −0.127, P = 0.59; n = 229). *PDCD1* expression also did not differ significantly between Black and White women in this cohort (β = −0.078, P = 0.63; n = 253). Expression varied less among patients in GSE268851 than in TCGA, with a standard deviation approximately 28% lower (**Fig. 2d**). Differences in tissue processing, RNA sequencing methods, and ancestry estimation may have contributed to the different findings, although the exact reasons remain unclear. Neither cohort showed a significant association between African ancestry and *CD274*, the gene encoding PD-L1 (TCGA P = 0.065; GSE268851 P = 0.19).

Overall, higher African ancestry was associated with increased *PDCD1* and several other immune-related gene-expression measures in TCGA, but the *PDCD1* association was not reproduced in GSE268851. Complete results and sensitivity analyses are provided in Supplementary Tables S2b, S2e, and S3.

### 3.3 Some negative TACSTD2–immune associations remain after epithelial adjustment

In GSE268851, higher *TACSTD2* expression was significantly associated with lower expression of five immune-related measures: *CD8A, GZMB, PRF1, PDCD1*, and the cytotoxic score (all P < 0.005; n = 229). Other epithelial markers, including *EPCAM, KRT8*, and *KRT18*, also showed mostly negative associations with these immune measures.

However, *TACSTD2* showed the strongest estimated negative associations with *CD8A, GZMB*, and the cytotoxic score (**Fig. 3a**).

**Fig. 3.**
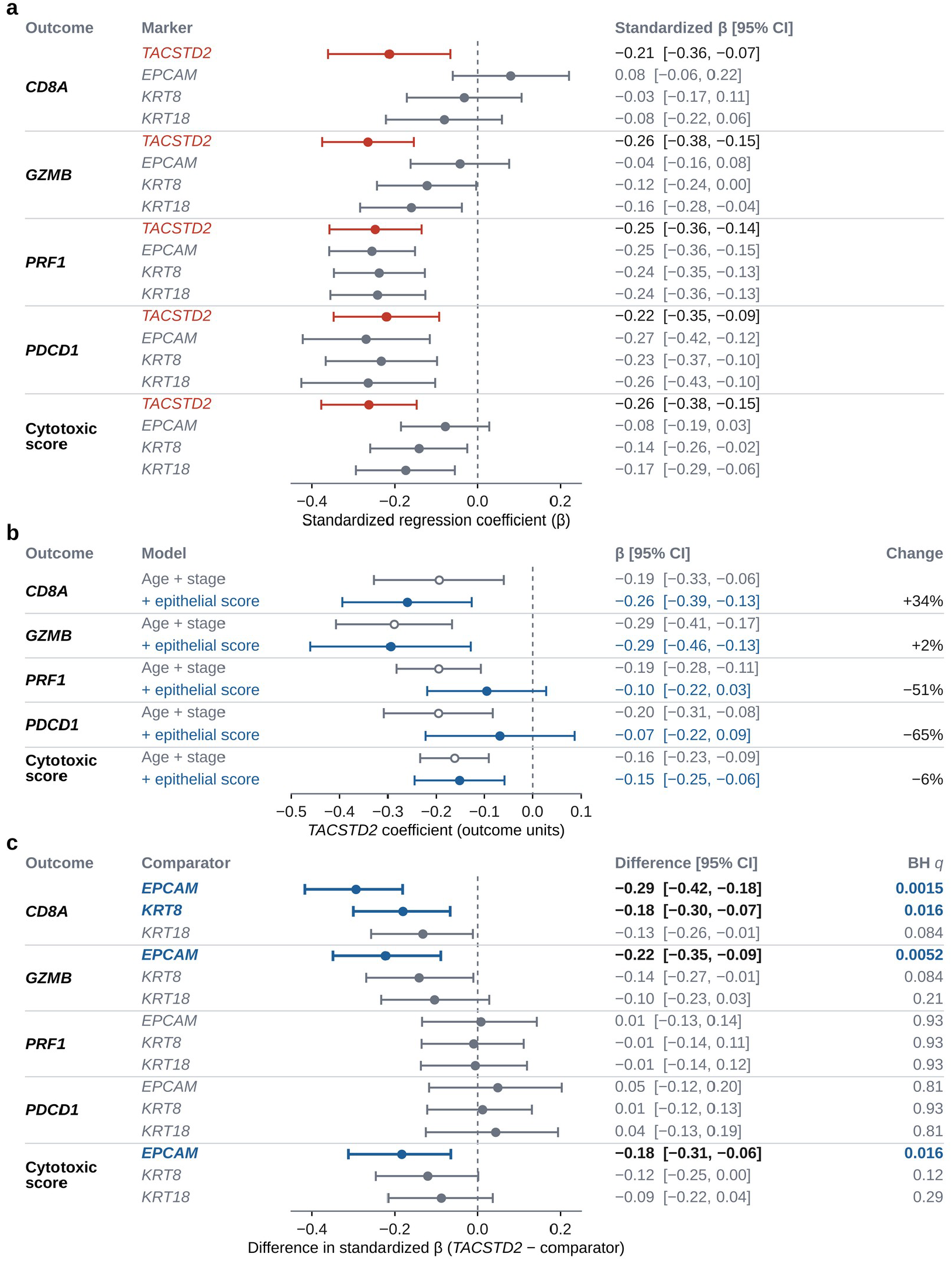
Associations of *TACSTD2* and other epithelial markers with immune-related gene expression in GSE268851 (n = 229). (a) Standardized regression coefficients for *TACSTD2, EPCAM, KRT8*, and *KRT18* with five immune-related measures, adjusted for age and tumor stage; *TACSTD2* is listed first within each outcome. (b) *TACSTD2* coefficients adjusted for age and stage (open circles) and after additional adjustment for the epithelial score (filled circles), which is based on *EPCAM, KRT8, KRT18*, and *CDH1*. Unstandardized coefficients are shown; Change gives the percentage change in absolute coefficient size after epithelial adjustment. (c) Differences in standardized coefficients between *TACSTD2* and each comparator marker (*TACSTD2* minus comparator), estimated from 10,000 paired patient-level bootstrap resamples; negative values indicate a stronger negative association for *TACSTD2*. Comparisons that remained significant after Benjamini–Hochberg correction (BH q < 0.05) are shown in bold. Points show coefficients or differences, and horizontal lines show 95% confidence intervals, which were not adjusted for multiple testing. Dashed lines mark zero

We next examined whether these relationships could be explained by differences in epithelial gene expression between tumor samples. After adjusting for an epithelial score based on *EPCAM, KRT8, KRT18*, and *CDH1*, the negative associations between *TACSTD2* and *PRF1* or *PDCD1* became weaker by 51% and 65%, respectively, and were no longer statistically significant (P = 0.13 and P = 0.39). In contrast, higher *TACSTD2* expression remained significantly associated with lower *CD8A* (P = 1.7 × 10^−4^), *GZMB* (P = 5.8 × 10^−4^), and cytotoxic score (P = 0.0016) after adjustment. All three associations remained significant after correction for multiple testing (q < 0.003; **Fig. 3b**). These findings show that differences in epithelial gene expression did not fully explain the negative associations with these three immune measures. In TCGA, none of the corresponding epithelial-marker associations remained significant after multiple-testing correction (Supplementary Tables S2c–S2d).

We then tested whether the negative associations observed for *TACSTD2* were statistically stronger than those of other epithelial markers. Four of the 15 comparisons remained significant after multiple-testing correction (**Fig. 3c**). Compared with *EPCAM, TACSTD2* showed significantly stronger negative associations with *CD8A* (coefficient difference = −0.294; q = 0.0015), *GZMB* (−0.222; q = 0.0052), and the cytotoxic score (−0.184; q = 0.016). Its negative association with *CD8A* was also significantly stronger than that of *KRT8* (−0.181; q = 0.016). No comparisons involving *KRT18, PRF1*, or *PDCD1* remained significant after correction.

Together, these results show that the negative relationships between *TACSTD2* and selected immune-related genes persisted after accounting for epithelial gene expression. In GSE268851, these associations were also significantly stronger than those of *EPCAM* for three immune measures and *KRT8* for *CD8A*.

### 3.4 TACSTD2 expression is higher in TNBC than normal breast and correlates with TROP2 protein abundance

*TACSTD2* expression was higher in TNBC tumors than in normal breast tissue, with median expression levels of 190.3 and 123.1 TPM, respectively. This represented a 1.55-fold difference (bootstrap 95% CI: 1.11–1.99; **Fig. 4a**). *PDCD1* expression was also 2.45-fold higher in TNBC tumors than in normal breast tissue (1.47 vs. 0.60 TPM). For *CD274*, the estimated difference was 1.35-fold, but the 95% confidence interval included 1, indicating greater uncertainty about this difference.

**Fig. 4.**
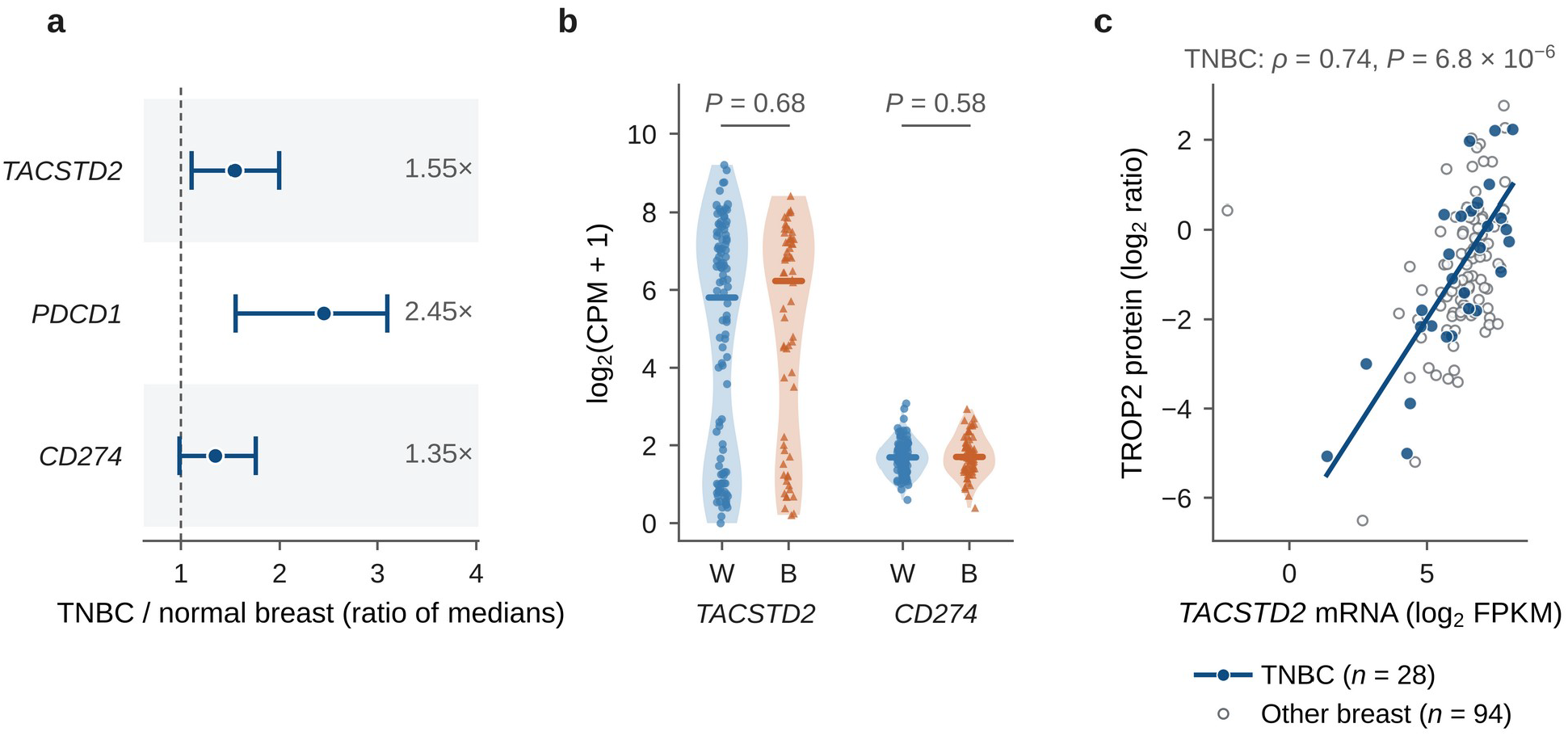
*TACSTD2* and immune-related gene expression in normal breast and TNBC, and the relationship between *TACSTD2* RNA and TROP2 protein. (a) Ratio of median expression in TCGA TNBC tumors (n = 107) to GTEx normal breast tissue (n = 80) for *TACSTD2, PDCD1*, and *CD274*. Points show ratios of medians, horizontal lines show bootstrap 95% confidence intervals, and the dashed line marks a ratio of 1. (b) *TACSTD2* and *CD274* expression in healthy breast tissue from White (W; circles; n = 108) and African American (B; triangles; n = 62) donors in GSE205725. Horizontal bars show group medians; P values are from Mann–Whitney tests. (c) *TACSTD2* mRNA expression (log2 FPKM) versus total TROP2 protein (log2 abundance ratio measured by mass spectrometry) in CPTAC breast tumors. Filled circles show TNBC tumors (n = 28), open circles show other breast tumors (n = 94), and the line shows a linear fit to TNBC tumors. Spearman ρ = 0.74 for TNBC and 0.56 for all breast tumors (n = 122)

We also examined whether these genes differed by race in healthy breast tissue. Neither *TACSTD2* nor *CD274* expression differed significantly between Black and White donors (P = 0.68 and P = 0.58, respectively; **Fig. 4b**). *PDCD1* was detectable in 94 of 170 donors but was generally expressed at low levels, with only five donors having expression above 1 CPM.

We next examined whether *TACSTD2* RNA expression was associated with TROP2 protein abundance. In 28 TNBC tumors from CPTAC, *TACSTD2* RNA expression showed a strong positive correlation with total TROP2 protein abundance (Spearman ρ = 0.74; 95% CI: 0.51–0.87; P = 6.8 × 10^−6^; **Fig. 4c**). A positive correlation was also observed across all 122 breast tumors (ρ = 0.56). These findings show that tumors with higher *TACSTD2* RNA expression generally had higher TROP2 protein abundance, although the analysis did not specifically measure TROP2 expression on the cell surface.

Finally, we examined five genes associated with response to SN-38, the cytotoxic drug delivered by sacituzumab govitecan. None showed significant associations with African genetic ancestry or race after correction for multiple testing (all q ≥ 0.25 across the three main analyses; Supplementary Figure S1 and Tables S4–S4b).

## 4. Discussion

We examined TROP2 and immune-related gene expression across racial and African ancestry groups with TNBC. Three findings stood out. *TACSTD2* was widely expressed without significant race or ancestry differences. The association between ancestry and *PDCD1* differed between cohorts. Some negative *TACSTD2*–immune associations remained after adjusting for epithelial gene expression and some were stronger than those for *EPCAM* and *KRT8*.

*TACSTD2* was detected in nearly all TNBC tumors and was higher in epithelial than immune regions, consistent with known TROP2 distribution [32, 33]. We found no significant race or ancestry differences. This supports studying TROP2-directed treatment across these populations, but does not prove equal cell-surface protein or treatment response. An exploratory ASCENT analysis reported benefit from sacituzumab govitecan in Black patients with metastatic TNBC, without examining ancestry [34]. TROP2 is also found in normal tissues, and its prognostic significance varies across cancers [33, 35].

In TCGA, tumors from patients with higher African ancestry showed higher *PDCD1* expression and several other T-cell-related measures, consistent with Martini et al. [3]. When we also adjusted for *CD3E*, a T-cell marker, the ancestry– *PDCD1* coefficient fell by about 65%. This suggests that differences in T-cell-related expression explain part of the association. However, we did not observe the same ancestry–*PDCD1* relationship in GSE268851. *PDCD1* also varied less among GSE268851 patients, which may have limited our ability to detect an association. Previous studies also disagree: O’Meara et al. found no consistent racial differences in the TNBC immune environment [16], while Yaghoobi et al. and Marczyk et al. reported greater immune infiltration or activity in African American TNBC [17, 18]. We therefore cannot describe higher *PDCD1* expression as a consistent feature of TNBC in women of African descent.

Previous studies found that TROP2-high breast tumors often show fewer immune-related features [19, 20]. Wu et al. provided experimental evidence that TROP2 can help keep immune cells out of tumors through a claudin-related mechanism [22]. Our study asked a more specific question: do the negative *TACSTD2*–immune relationships remain when we account for epithelial gene expression, and are they stronger than those involving other epithelial markers?

In GSE268851, the negative *TACSTD2* associations with *CD8A, GZMB*, and the cytotoxic score remained significant after adjusting for epithelial gene expression. The relationships with *PDCD1* and *PRF1* became weaker and were no longer significant. This means that differences in epithelial expression did not fully explain every *TACSTD2*–immune association. In direct statistical comparisons, *TACSTD2* was more negatively associated than *EPCAM* with *CD8A, GZMB*, and the cytotoxic score, and more negatively associated than *KRT8* with *CD8A*. It did not differ significantly from *KRT18*. These findings are consistent with an immune-poor pattern in tumors with high TROP2 expression, but our bulk RNA data cannot show that TROP2 itself causes immune cells to be excluded.

*TACSTD2* and *PDCD1* expression were 1.55-fold and 2.45-fold higher, respectively, in TNBC tumors than in normal breast tissue. In healthy breast samples, neither *TACSTD2* nor *CD274* differed significantly between Black and White donors. *TACSTD2* RNA also correlated strongly with total TROP2 protein in CPTAC TNBC tumors. This supports the usefulness of RNA data for studying TROP2 abundance, but not for measuring how much TROP2 is available on the tumor-cell surface. That distinction matters because antibody-drug conjugates must reach and bind their target on cancer cells. TROP2 is also present in normal breast tissue [33].

We also tested five genes related to SN-38 activity or drug transport because SN-38 is delivered by sacituzumab govitecan. None showed significant race or ancestry associations after multiple-testing correction. Previous work connects *SLFN11* and ABC transporters to treatment response [36, 37]. However, these results do not prove equal drug sensitivity across populations. Drug metabolism, including *UGT1A1*-related toxicity, intracellular processing, and payload sensitivity also matter [38–41]. Variation in TROP2 staining tests and preclinical responses supports protein-level validation [42, 43].

Our study has limitations. The negative associations between *TACSTD2* and immune-related genes were clearer in GSE268851. In TCGA, the corresponding associations were not significant after correction for multiple testing. Cohorts differed in participants, tissue processing, RNA sequencing, and ancestry estimation. Our epithelial score is based on gene expression, not measured tumor purity. Bulk RNA cannot tell whether lower immune-gene expression reflects fewer immune cells or changes in their activity. Tumor and normal samples were not matched, and treatment response was not assessed. Independent cohorts, spatial or single-cell studies, and protein measurements can test these findings further.

Overall, *TACSTD2* was widely expressed in the TNBC populations examined, while the ancestry–*PDCD1* relationship differed between cohorts. In GSE268851, some negative *TACSTD2*–immune associations remained after epithelial adjustment and were stronger than those for *EPCAM* and, for *CD8A, KRT8*. These results identify specific relationships that should be tested in independent biological and clinical studies.

## Abbreviations

ADC: antibody-drug conjugate
AFR: African ancestry proportion
BH: Benjamini–Hochberg
BMI: body mass index
CI: confidence interval
CPM: counts per million
CPS: combined positive score
CPTAC: Clinical Proteomic Tumor Analysis Consortium
ER: estrogen receptor
FFPE: formalin-fixed paraffin-embedded
FPKM: fragments per kilobase of transcript per million mapped reads
GTEx: Genotype-Tissue Expression
HC3: heteroskedasticity-consistent covariance estimator type 3
HER2: human epidermal growth factor receptor 2
IHC: immunohistochemistry
OLS: ordinary least squares
PD-1: programmed cell death protein 1
PD-L1: programmed death-ligand 1
PR: progesterone receptor
SD: standard deviation
TCGA: The Cancer Genome Atlas
TNBC: triple-negative breast cancer
TPM: transcripts per million
TROP2: trophoblast cell-surface antigen 2.

## Statements and Declarations

### Ethics approval

This study used publicly available, de-identified datasets and involved no new recruitment or specimen collection; therefore, institutional review board approval was not required.

### Consent to participate

Not applicable.

### Consent to publish

Not applicable.

### Data availability

GSE268851 and GSE205725 are available from GEO; TCGA-BRCA from the Genomic Data Commons; Toil expression from UCSC Xena; and CPTAC from cBioPortal (brca_cptac_2020). The Zhu spatial dataset is available at https://doi.org/10.5281/zenodo.12752405. Ancestry estimates were obtained from the source-study supplements and Carrot-Zhang et al. [4, 23]. Processed model results, sample summaries, and supporting tables are included in the supplementary workbook.

### Code availability

Software versions, input checksums, and detailed reproducibility information are provided in the supplementary files. Analysis scripts are available from the authors upon reasonable request.

## Acknowledgments

The authors acknowledge the BigCARE cancer research training program at the University of California, Irvine, for bioinformatics training and research support. Computational analyses used the Anvil supercomputer at Purdue University, made available through the Advanced Cyberinfrastructure Coordination Ecosystem: Services & Support (ACCESS) program. Anvil is supported by the U.S. National Science Foundation (award OAC-2005632). The ACCESS allocation identifier is TRA220018.

## Funding

No specific funding supported this study.

### Author contributions

A.O. conceived the study, assembled the datasets, developed the analyses and figures, and drafted the manuscript. E.O.A. supervised the work and contributed to study design, interpretation and manuscript revision. All authors are responsible for the final content.

### Supplementary Information

The SN-38 gene panel is retained in supplementary Tables S4–S4b; Supplementary Figure S1 is supplied as a separate supplementary figure file. The workbook contains a Contents sheet with descriptions of each table. S1 maps applicable REMARK items; S2 and S2b-S2f contain current target models, PDCD1 sensitivities, marker models, epithelial adjustment, jackknife results and paired contrasts. S2g-S2i provide patient inclusion lists and all coefficients for key models. S3 reports 19 immune-panel tests. S4-S4b contain detection and five-gene payload results, including within-Black analyses. S5-S6 report BMI and recorded-chemotherapy sensitivities. S7-S8 identify inputs and software; S9-S11 contain expression and protein observations and summaries. S12 summarizes baseline characteristics. Current results are also supplied as TSV files.

