## Supplementary Figure S1 for "TROP2 and immune gene expression by African ancestry in triple-negative breast cancer"

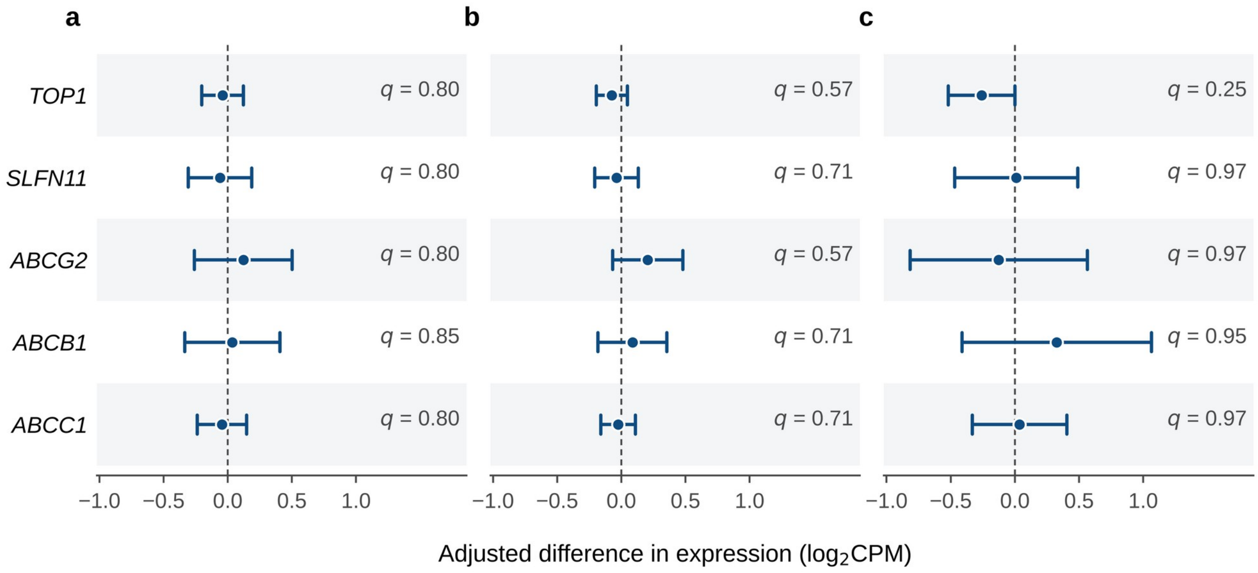


**Supplementary Fig. S1** SN-38-related gene expression by African ancestry and race in TNBC. Adjusted differences in expression of *TOP1*, *SLFN11*, *ABCG2*, *ABCB1*, and *ABCC1* (a) per unit of African ancestry in GSE268851 (n = 229), (b) between Black and White women in GSE268851 (n = 253), and (c) per unit of African ancestry in TCGA (n = 107). Models were adjusted for age, tumor stage, and epithelial score. Points show regression estimates, horizontal lines show 95% confidence intervals, and dashed lines mark no difference. q values are Benjamini–Hochberg-adjusted across the five genes within each analysis. *UGT1A1* was excluded before analysis because of insufficient detection
